# Closing the loop between brain and electrical stimulation: A proof-of-concept randomized trial of real-time fMRI-guided tACS optimization

**DOI:** 10.64898/2026.02.28.708766

**Authors:** Ghazaleh Soleimani, Rayus Kuplicki, Beni Mulyana, Aki Tsuchiyagaito, Masaya Misaki, Martin P Paulus, Hamed Ekhtiair

**Author notes:** **Corresponding Author** Hamed Ekhtiari, MD, PhD Department of Psychiatry, University of Texas Southwestern (UTSW), Dallas, TX.

## Abstract

Closed-loop transcranial alternating current stimulation (tACS) combined with real-time functional MRI (tACS–fMRI) enables adaptive optimization of noninvasive brain stimulation based on ongoing neural dynamics. We implemented one of the first randomized, double-arm, closed-loop tACS–fMRI protocols to iteratively adjust stimulation frequency and phase alignment between the right dorsolateral prefrontal cortex (F4) and right inferior parietal cortex (P4) to modulate frontoparietal functional connectivity (FFC) during a 2-back working memory task. Twenty healthy adults were randomized to an up-regulation group (n = 10), receiving parameters optimized to maintain or enhance FFC, or a down-regulation group (n = 10), receiving parameters optimized to reduce FFC using a simplex-based adaptive algorithm across two training runs. Individualized optimized parameters were applied during a subsequent test run, with resting-state fMRI acquired before and after stimulation. During the test phase, connectivity trajectories diverged across groups: FFC was preserved in the up-regulation group but declined in the down-regulation group (signed-rank p = 0.019), with a significant between-group difference confirmed by permutation testing (20,000 iterations; p = 0.043). Behavioral effects were primarily observed in the accuracy learning dynamics. Although overall mean performance was comparable, the up-regulation group demonstrated a more positive accuracy trajectory, greater within-test accuracy gain (p = 0.036), and steeper trial-wise improvement (p = 0.035). Reaction time decreased across runs in both groups, consistent with practice effects, with no significant group differences in gain; however, the test-run RT slope showed a trend toward a steeper reduction in reaction time in the up-regulation group relative to the down-regulation group (p = 0.065). Resting-state analyses revealed significant time-by-group interaction clusters in seed-based connectivity (voxel-wise p < 0.001; cluster-level FDR p < 0.05), with connectivity increases observed in the up-regulation group relative to the down-regulation group, indicating persistent network modulation following stimulation. These findings demonstrate the feasibility of closed-loop tACS–fMRI for individualized network-targeted neuromodulation and suggest that real-time connectivity optimization selectively stabilizes task-relevant connectivity and enhances accuracy learning while producing sustained effects on intrinsic brain networks.

## Introduction

Working memory is a cognitive function responsible for temporarily holding information needed to perform complex tasks such as reasoning, comprehension, and learning ^1^. The connection between the dorsolateral prefrontal cortex (DLPFC) and posterior parietal cortex (PPC) forms the frontoparietal network (FPN), a dynamic system involved in regulating executive functions such as working memory ^2^. It is hypothesized that communication within and across the FPN is mediated by oscillatory synchronization of neuronal activity within specific phase and frequency ranges ^3,4^. This rhythmic coordination is thought to strengthen FPN, and both animal and human studies confirmed that higher frontoparietal functional connectivity (FFC) following working memory training was positively correlated with higher working memory performance^5-7^.

This FFC can be modulated via exogenous stimulation. For example, transcranial alternating current stimulation (tACS) is a non-invasive neuromodulation technique that delivers sinusoidal electrical currents to modulate endogenous neural oscillations at specific frequencies ^8^. By entraining ongoing rhythmic activity, tACS can influence interregional synchronization and thereby modulate large-scale network communication during cognitively demanding processes such as working memory^9^. High definition dual-site tACS extends this framework by concurrently stimulating two anatomically and functionally connected cortical regions, like DLPFC and PPC, as main nodes of FPN^10^. However, regarding the nature of brain traveling waves, stimulation effects depend not only on the applied frequency but also critically on the relative phase offset between the two sites ^11^.

Electrophysiological and computational evidence indicate that optimal phase and frequency vary substantially across individuals and may fluctuate within individuals over time^12,13^. Such inter- and intra-individual variability challenges the assumption that fixed stimulation protocols effectively align with endogenous network dynamics. Despite this, based on our recent systematic review ^14-16^, most prior dual-site tACS studies targeting frontoparietal synchronization have employed standardized, one-size-fits-all stimulation parameters across participants, potentially limiting the consistency and magnitude of network-level and behavioral effects ^17,18^.

Closed-loop neuromodulation approaches using real-time neuroimaging (e.g., EEG or fMRI) offer a promising solution by enabling dynamic assessment of brain function and online optimization of stimulation parameters ^19,20^. Unlike traditional open-loop paradigms that apply fixed parameters across participants, closed-loop systems can continuously adapt stimulation to an individual’s ongoing neural state, thereby accounting for both inter- and intra-individual variability in oscillatory dynamics ^21^. Previous closed-loop tACS–EEG studies suggest that stimulation outcomes depend on both phase and frequency, with promising effects on sleep-related memory consolidation and α-rhythm modulation ^22-24^. A key limitation, however, is that concurrent tACS–EEG is heavily contaminated by artifacts, making it unsuitable for real-time parameter adjustment ^25^. To address this, for example, intermittent protocols were used, delivering short stimulation epochs separated by brief EEG recordings to recalibrate frequency^26^. In parallel, fMRI-guided approaches, with an established concurrent tES-fMRI checklist ^27^, offer a complementary solution, enabling dynamic optimization of stimulation while providing causal insights into how brain networks respond in real time.

We previously introduced a conceptual framework for closed-loop tES-fMRI protocols and registered a proof-of-concept study demonstrating the feasibility and signal-to-noise characteristics of the system ^28,29^. Building on this foundation, the present study employed a randomized, double-arm, closed-loop, online tACS-fMRI design to optimize stimulation parameters, specifically phase and frequency, for enhancing working memory performance in healthy participants via FFC. The primary objectives were: (1) to determine whether real-time closed-loop optimization could identify stimulation parameters that enhance (up regulation) or disrupt (down regulation) the target FFC during training runs; (2) to evaluate whether these optimized parameters improve online FFC and performance during a subsequent working memory testing run; and (3) to examine changes in offline resting-state functional connectivity before and after phase- and frequency-adjusted stimulation across up- and down-regulation groups.

## Method

### Participants

Thirty-three healthy young adults were recruited through the Laureate Institute for Brain Research. Inclusion criteria required participants to be between 18 and 60 years of age, have no history of neurological disorders. All participants provided written informed consent before the start of the study. The study protocol was approved by the Western Institutional Review Board (WIRB) #20200192 and conducted in accordance with the Declaration of Helsinki. Of the 33 individuals assessed for eligibility, 10 were excluded (1 declined to participate, and 9 were used for protocol development). The remaining 23 participants were randomly assigned to either the up-regulation group (n = 11) or the down-regulation group (n = 12). In the up-regulation group, one participant withdrew due to scalp pain during stimulation, resulting in 10 participants included in the final analysis. In the down-regulation group, one participant was unable to begin stimulation due to high impedance, and another could not complete the session due to connection issues. Of the remaining 10 participants, one was excluded from analysis due to excessive censored time points (number of TRs exceeding the censoring threshold), leaving 9 participants in the final analysis for the down-regulation group (see the consort diagram in Supplementary Section S1).

### Study Design

Participants were randomly assigned to one of two groups: an up-regulation group (max-optimized) or a down-regulation group (min-optimized). Both groups completed two training runs while receiving tACS combined with real-time fMRI. In the up-regulation group, an adaptive algorithm searched for the stimulation parameters (frequency and phase difference) that maximized functional connectivity between the right dorsolateral prefrontal cortex (F4) and the right inferior parietal cortex (P4). In the down-regulation group, the same procedure was followed, but the algorithm aimed to find parameters that minimized this connectivity.

Each training part consisted of 15 stimulation blocks (20 seconds of stimulation followed by 10 seconds of no stimulation). Following the training runs, participants underwent a testing run where stimulation parameters were fixed: the up-regulation group received the previously optimized parameters, while the down-regulation group received the min-optimized parameters identified during training. A resting-state fMRI scan (6 minutes and 50 seconds) was conducted before and after the optimization parts and again after the testing run. A 7-minute washout period, including a resting-state scan, was included between training and testing parts to reduce potential aftereffects of stimulation. To maintain consistent cognitive engagement and assess behavioral outcomes, participants performed a 2-back working memory task throughout the training and testing runs (more technical details can be found here^28^).

### Concurrent tACS-fMRI Data Collection

Concurrent tACS-fMRI data were collected while participants lay supine inside the MRI scanner, fitted with 10 high-definition (HD) electrodes arranged in a frontoparietal montage. The tACS device was positioned outside the MRI room and connected to the participant via an RF filter box mounted on the MRI penetration panel based on the checklist developed by our team to run a concurrent tACS-fMRI ^27^. Stimulation signals were delivered through shielded cables to minimize electromagnetic interference. An MR-compatible stimulator cable connected the tACS device to the filter box, and stimulation was applied through banana cables routed to the participant’s scalp. All components were secured and arranged to comply with MRI safety standards and minimize artifacts. The entire setup was optimized for simultaneous stimulation and image acquisition, ensuring low artifact levels and preserving signal-to-noise ratio. Further technical details regarding MRI safety and signal quality under stimulation can be found in our previous publication ^28^.

### Working Memory Task

A letter-based 2-back task was used to assess the effects of stimulation conditions on working memory performance ^30^. Before the first stimulation session, participants completed a practice block to become familiar with the task. During each stimulation session, participants performed the working memory task under either up-regulation or down-regulation stimulation. Each session consisted of 112 trials, during which participants were instructed to respond within 2 seconds after stimulus onset. Behavioral responses were recorded using a response box, capturing both reaction time (RT, defined as the time from stimulus onset to response) and accuracy (percentage of correctly identified targets (hits) and correctly rejected non-targets compared to the total number of trials).

### Stimulation Setup

Two 4×1 MR-compatible high-definition (HD) electrode montages were used to target the right dorsolateral prefrontal cortex (centered at F4) and the right inferior parietal cortex (centered at P4). Each stimulation site received 1 mA peak-to-peak current at the central electrode (F4 or P4), with 0.25 mA return currents delivered to the surrounding electrodes, positioned 180° out of phase. In each stimulation site, return electrodes were positioned at an equal distance of 3 cm from a central electrode. The right frontoparietal network was selected based on its well-established role in supporting working memory, with evidence suggesting that the right hemisphere frontoparietal system exerts stronger effects on executive functions ^31^. The stimulation frequency and phase offset between the F4 and P4 electrodes were optimized in each session according to the intrinsic connectivity profile of the frontoparietal network. The stimulation setup was similar in both groups.

### Optimization Algorithm

During tACS-fMRI, a Nelder–Mead Simplex optimization algorithm ^32^ was implemented during two training phases to identify tACS parameters (frequency and phase difference) that either maximize or minimize FFC ^28^. The optimization was conducted in real-time, adapting stimulation parameters based on the observed fMRI connectivity between frontal and parietal sites. We used AFNI, in-house Python scripts, and MATLAB for the online FFC calculation and optimization ^33^. Each subject completed 30 blocks across Training 1 and Training 2. Each block consisted of 20 seconds of tACS using frequency and phase values chosen by the Simplex optimizer, followed by 10 seconds of rest. During each block, online FFC was calculated in real time with a 20-second sliding window (10 TRs). We normalized the distribution using Fisher’s z-transformation. The FFC z values were used as online FFC values and were then used by the Simplex algorithm to determine the parameters for the next block. In this process, a block was considered successful if the parameters proposed by the Simplex rules (reflection, expansion, contraction, or shrink) were accepted and incorporated as a new vertex in the parameter space, thereby advancing the optimization. A block was considered unsuccessful if the proposed parameters were rejected by the algorithm and did not contribute to updating the optimization path. For the Simplex optimizer, the initial search triangle in the theta band was defined by the vertices (frequency in Hz, phase difference in degrees): [6, 5], [10, −3], and [2, −3]. This configuration accelerates convergence by centering the search around 6 Hz and 0°, values suggested by prior literature as optimal starting points.

The Simplex method begins with an initial triangle in parameter space. At each iteration, it identifies the worst vertex and replaces it with a new point through reflection, expansion, contraction, or shrink steps, guided by the positions of the vertices. For the up-regulation group, the goal was to identify parameters that maximize FFC, while the down-regulation group followed a mirrored procedure aiming to minimize FFC. At the end of training, the best-performing parameter set was selected based on the highest (for up-regulation) or lowest (for down-regulation) observed online FFC.

### Behavioral Data Analysis

Behavioral performance during the 2-back working memory task was quantified using trial-level accuracy (correct/incorrect responses) and reaction time (RT, in seconds). Analyses focused on the Test run, which followed two training runs (TR1 and TR2). Trials with missing responses or RT values ≤ 50 ms were excluded. For each participant, overall Test performance was indexed by mean RT across valid Test trials and by the proportion of correct responses. These measures provided summary indices of steady-state task performance during the Test phase.

To characterize within-run performance dynamics relevant to the iteration-level optimization framework, gain measures were computed separately for RT and accuracy. The 120 Test trials were partitioned into early (trials 1–40) and late (trials 81–120) segments to capture session-level adaptation. RT gain was defined as the difference between early and late RT (RT_gain = RT_early − RT_late), such that positive values indicate faster responses by the end of the run. Accuracy gain was defined as the difference between late and early accuracy (Accuracy_gain = Accuracy_late − Accuracy_early), with positive values reflecting improved accuracy over time. This approach enabled dissociation between overall performance level and within-session adaptation, allowing evaluation of whether stimulation influenced steady-state performance, learning dynamics, or both.

To further quantify learning trajectories, slope-based metrics were derived for both RT and accuracy using two complementary approaches. First, a trajectory slope was estimated to capture coarse learning trends across the run. Trials were divided into six equal bins (20 trials per bin), and mean performance within each bin was computed. For each participant, a linear regression was fitted to these bin means as a function of trial position, yielding a trajectory slope that summarizes directional performance change while reducing trial-level variability.

Second, a Test-phase slope was calculated using all individual Test trials to provide a fine-grained estimate of learning dynamics. For RT, slopes were obtained from linear regression of log-transformed RT on trial index to account for the positively skewed RT distribution. For accuracy, slopes were estimated from linear regression of trial-wise accuracy on trial index. Negative RT slopes indicate progressive speeding across trials, whereas positive accuracy slopes indicate improved accuracy over time. These slope measures complement gain indices by capturing continuous learning trends rather than discrete early–late changes.

Between-group differences (Up-regulation vs. Down-regulation) in Test mean RT, Test mean accuracy, RT gain, accuracy gain, and slope measures were evaluated using two-tailed Wilcoxon rank-sum tests (Mann–Whitney U tests), given the modest sample size and potential non-normality of behavioral distributions. Descriptive statistics are reported as mean ± standard error of the mean (SEM) and medians where appropriate. All statistical tests were two-tailed with a significance threshold of α = 0.05.

### Task-based fMRI analysis

Functional connectivity was quantified across 15 predefined connections for each participant during three sequential runs (TR1, TR2, and Test). Participants were a priori assigned to the up-regulation or down-regulation group based on study ID. For each participant and run, connectivity values were first organized at the connection level and then summarized at the subject level by computing the mean connectivity across the 15 connections. These subject-level mean values were used to characterize overall connectivity per run. To examine within-session connectivity dynamics during the Test run, we additionally computed a gain measure defined as the difference between late and early connectivity within Test (mean of connections 11–15 minus mean of connections 1–5). Positive values indicate increasing connectivity across the run. For exploratory analyses of within-run trajectory, linear slopes were estimated for each participant by fitting connectivity as a function of iteration number (1–15). The primary outcome was the change in connectivity from Training to Test. Training connectivity was defined as the mean of TR1 and TR2 connectivity, and Test change (ΔTest) was computed as Test minus Training mean for each participant. Normality of connectivity distributions within each group and run was assessed using the Lilliefors (Kolmogorov–Smirnov) test. Given the modest sample size and evidence of non-normality in several conditions, nonparametric tests were used as primary inferential procedures.

Within-group changes (TR1 vs. Test and TR2 vs. Test) were evaluated using Wilcoxon signed-rank tests. Between-group differences in ΔTest, Test mean connectivity, and Test gain were assessed using two-sided Wilcoxon rank-sum tests. For completeness, Welch’s two-sample t-tests (assuming unequal variances) were also conducted, and results were considered convergent when both parametric and nonparametric tests yielded consistent conclusions. As an additional robustness analysis, a two-sided permutation test (20,000 label shuffles) was performed on the absolute group mean difference in ΔTest to obtain an empirical p-value independent of distributional assumptions. All statistical tests were two-tailed with a significance threshold of α = 0.05.

### Resting state fMRI analysis

For each subject, the structural T1-weighted MRI was used to construct computational head models. The peak electric field (99th percentile) within the frontal and parietal cortices was identified and mapped to MNI space. A 10-mm sphere was centered on each peak location and intersected with an MNI brain mask to exclude non-brain and white matter voxels. Seed-to-whole-brain resting-state functional connectivity was computed using the CONN toolbox (22.v2407)^34^. Analyses were performed separately for the frontal and parietal seeds across each resting-state run (rest-fMRI 1, 2, and 3; Figure 1). A time-by-group interaction analysis was subsequently performed to assess whether stimulation altered functional connectivity in response to individualized tACS. A voxel-wise threshold of p < 0.001 (uncorrected) and a cluster-level threshold of p < 0.05 (FDR-corrected) were applied. To characterize the anatomical distribution of significant clusters, each cluster was intersected with the 7-network parcellation of Yeo et al. (2011). The number of voxels overlapping with each large-scale network was computed, and the dominant network coverage is reported in Table 2.

**Figure 1.**
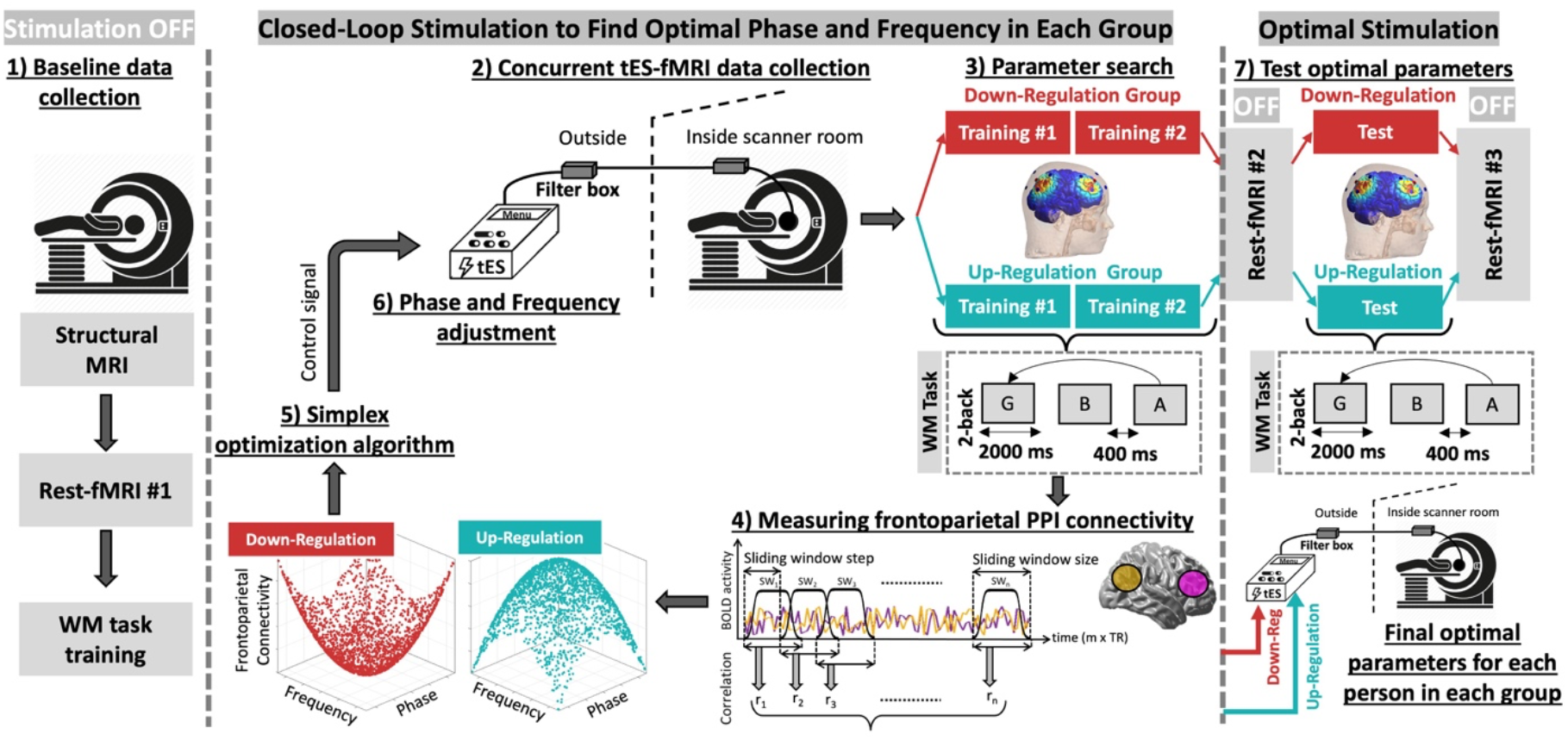
Study Design for Concurrent tACS-fMRI Optimization Protocol. Participants were randomly assigned to either an up-regulation group (max-optimized) or a down-regulation group (min-optimized). The study commenced with the collection of baseline data, which included structural MRI, resting-state fMRI, and a brief working memory (WM) task training (Panel 1). Participants then underwent two training runs of concurrent transcranial alternating current stimulation (tACS) and real-time fMRI (Panel 2), during which an adaptive simplex optimization algorithm adjusted the stimulation phase and frequency to modulate frontoparietal functional connectivity between the right dorsolateral prefrontal cortex (F4) and right inferior parietal cortex (P4) (Panel 3). In the up-regulation group (blue), the algorithm aimed to maximize connectivity; in the down-regulation group (red), it aimed to minimize it. Each training run consisted of 15 blocks of stimulation (20 seconds on, 10 seconds off, TR = 2), followed by a 7-minute washout period with resting-state fMRI. In the subsequent testing phase (Panel 4), participants received stimulation with the final optimized parameters identified during training (either maximizing or minimizing connectivity, depending on the group) while performing a 2-back WM task. Resting-state fMRI scans were collected before and after the training and testing sessions to assess changes in intrinsic connectivity patterns.

**Figure 2.**
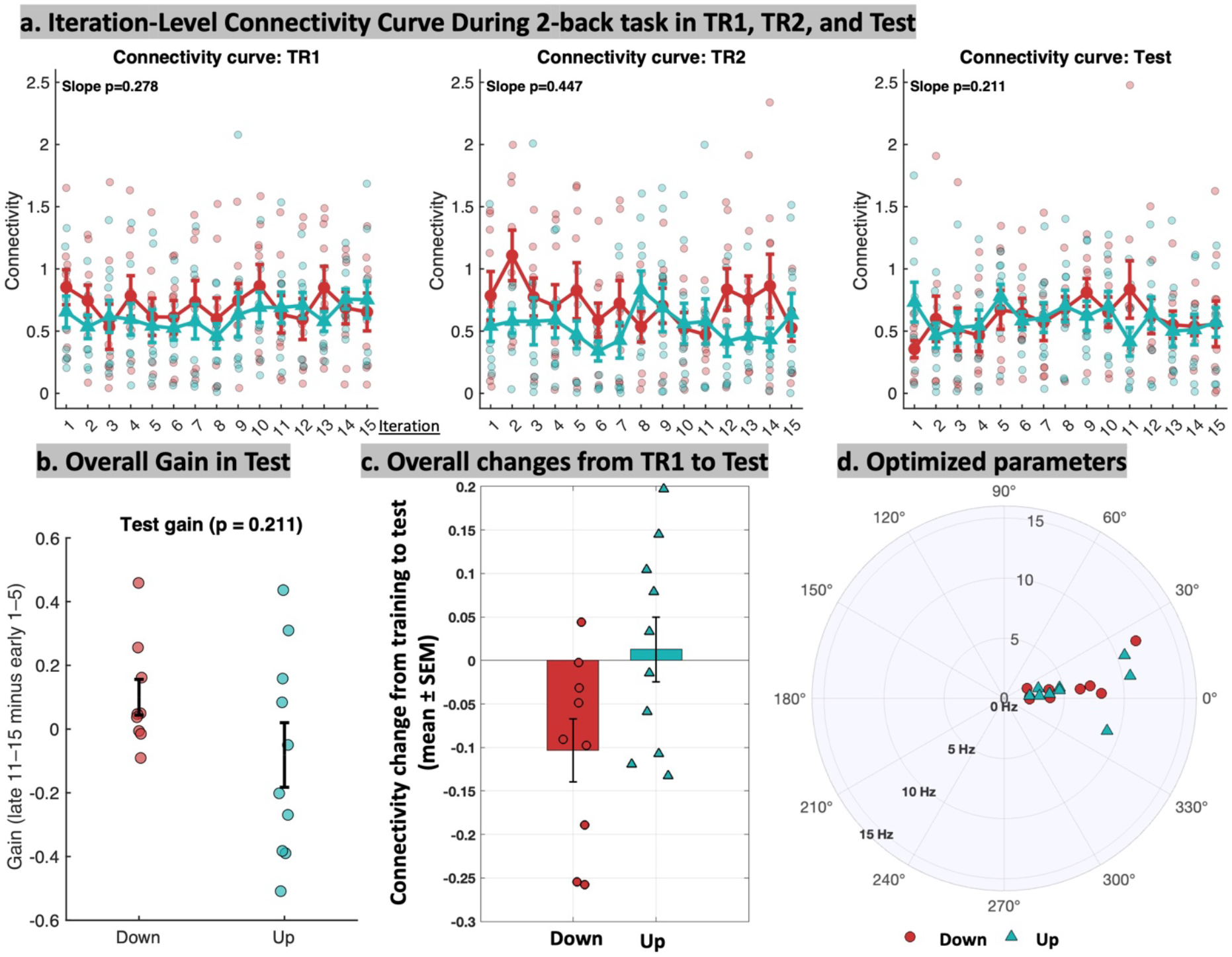
Iteration-level functional connectivity dynamics and optimized stimulation parameters during the 2-back task. (a) Iteration-level functional connectivity curves during TR1, TR2, and Test. Points represent individual participants at each iteration (1–15), color-coded by group (Down-regulation in red; Up-regulation in teal). Solid lines with error bars denote group mean ± standard error of the mean (SEM). P-values indicate between-group differences in within-run connectivity slope (linear fit across iterations). (b) Overall connectivity gain during the Test run, defined as the difference between late (iterations 11–15) and early (iterations 1–5) connectivity. Points represent individual participants; error bars indicate mean ± SEM. (c) Overall change in connectivity from training to Test, computed as the difference between Test mean connectivity and mean connectivity across TR1 and TR2. Bars represent group mean ± SEM with individual subject values overlaid. (d) Final optimized stimulation parameters for each participant displayed in polar coordinates. Radial distance represents stimulation frequency (Hz), and angular position represents phase difference (degrees). Each point corresponds to one participant, color-coded by group.

### Blinding and tolerability

After obtaining written informed consent, participants were randomized during the baseline session to receive either up-regulated or down-regulated stimulation parameters using a computer-generated allocation code. This procedure ensured that participants were blinded to the stimulation condition. Blinding efficacy was assessed at the end of the stimulation session by asking participants to indicate which group they believed they had been assigned to (blinding guess and confidence in guess). Side effects were evaluated at the end of the session using a standard transcranial electrical stimulation side-effect questionnaire, and responses were compared between groups.

## Results

The CONSORT diagram outlining participant inclusion and group allocation is presented in Figure S1. Two groups are well-matched in terms of demographic characteristics (Table 1).

**Table 1.** Demographic data, blindness, and side effects of the stimulation.

| | Up-Regulation Group<br>(n = 10) Mean ± SD | Down-Regulation Group<br>(n = 10) Mean ± SD | Statistic |
| --- | --- | --- | --- |
| **Age** | 38.2 ± 10.35 | 38.6 ± 10.46 | t(18) = -0.09, p = 0.93 |
| **Male (%)** | 4 (40%) | 4 (40%) | $\chi^2(1) < 0.001$, p = 1.00 |
| **POMS** |  |  |  |
| Total Mood Disturbance | -7.20 ± 17.37 | -6.70 ± 9.62 | t(18) = -0.08, p = 0.94 |
| Tension | 3.40 ± 2.99 | 2.50 ± 1.90 | t(18) = 0.80, p = 0.43 |
| Depression | 2.10 ± 4.98 | 0.20 ± 0.42 | t(9.13) = 1.20 p = 0.26 |
| Anger | 1.10 ± 2.85 | 0.70 ± 1.57 | t(18) = 0.39, p = 0.70 |
| Fatigue | 2.00 ± 2.16 | 1.40 ± 1.96 | t(18) = 0.65, p = 0.52 |
| Vigor | 18.70 ± 6.11 | 14.10 ± 7.00 | t(18) = 1.57, p = 0.13 |
| **STAI-State** | 29.10 ± 6.84 | 29.90 ± 7.17 | t(18) = -0.26, p = 0.80 |
| **Blinding** |  |  |  |
| Blinding guess | 5 (50%) | 6 (60%) | $\chi^2(1) = 3.2$, p = 0.07 |
| Confidence in guess (0-10) | 4 | 6 | t(18) = 0.48, p = 0.64 |
| **Side Effects** |  |  |  |
| Headache | 1 (10%) | 0 | $\chi^2(1) = 1.05, p = 0.30$ |
| Scalp pain | 2 (20%) | 3 (30%) | $\chi^2(1) = 0.27, p = 0.60$ |
| Tingling | 3 (30%) | 3 (30%) | $\chi^2(1) = 0.0, p = 1$ |
| Itching | 1 (10%) | 0 | $\chi^2(1) = 1.05, p = 0.3$ |
| Burning sensation | 1 (10%) | 2 (20%) | $\chi^2(1) = 0.39, p = 0.5$ |
| Skin redness | 1 (10%) | 0 | $\chi^2(1) = 1.05, p = 0.3$ |
| Sleepiness | 5 (50%) | 6 (60%) | $\chi^2(1) = 0.20, p = 0.65$ |
| Neck pain | 1 (10%) | 1 (10%) | $\chi^2(1) = 0.0, p = 1$ |
| Trouble concentrating | 4 (40%) | 2 (20%) | $\chi^2(1) = 0.95, p = 0.33$ |
| Acute mood change | 0 | 0 | $\chi^2(1) = 0.0, p = 1$ |

POMS: Profile of Mood Stats. Results are reported in mean standard deviation (SD), t(df): t-statistic, p: uncorrected p-value, $\chi^2$ : Chi-square statistic, Welch's t for Depression score: A variant of the t-test used when group variances are unequal. Blindness: number of participants who correctly identified their assigned group. Blindness and side effects results are reported in the number of subjects and their percentage, t(df): t-statistic, p: uncorrected p-value, $\chi^2$ : Chi-square statistic.

**Table 2.** Significant resting state clusters connected to the frontal seed region. Seed-to-whole-brain functional connectivity was computed from individualized frontal target (defined by peak electric field locations) across Rest3 vs Rest2 (as described in Figure 1, before and after application of the final optimized stimulation). The table reports clusters showing significant time-by-group interactions, with voxel-wise threshold set at *p* < 0.001 (uncorrected) and cluster-level FDR-corrected *p* < 0.05.

| [x, y, z] MNI coordinate of Center of Mass (COM) | Size | FDR-corrected p | Main covered large-scale network |  |  |  |  |  |  |
| --- | --- | --- | --- | --- | --- | --- | --- | --- | --- |
|  |  |  | FPN | DMN | Lim | Vis | SoM | SN | DAN |
| [12, -12, -28]-Cerebellum | 4454 | 0.00003 | 391 | 212 | 219 | 1109 | 276 | 171 | 31 |
| [42, 36, -8]-MFG | 1497 | 0.04670 | 332 | 276 | 2 | 0 | 1 | 69 | 1 |
| Total | 5951 |  | 723 | 488 | 221 | 1109 | 277 | 240 | 32 |
|  |  |  | 12% | 8% | 3% | 19% | 5% | 4% | 0.5% |

The coverage of the clusters within the seven large-scale resting-state networks in the Yeo atlas is reported. **Abbreviations:** FPN, frontoparietal network; DMN, default mode network; Lim, limbic network; Vis, visual network; SoM, somatomotor network; SN, salience network; and DAN, dorsal attention network.

Blinding was largely successful, with no significant difference between groups in participants’ ability to correctly guess their stimulation condition (χ^2^(1)=3.2, p=0.07) or in confidence ratings of their guesses (t(18)=0.48, p=0.64). Reported side effects were mild and infrequent, and none differed significantly between the groups (all p>0.30). The most commonly reported effect was sleepiness (50–60%), followed by tingling and difficulty concentrating, while no acute mood changes or other adverse effects were reported in either group (Supplementary Section S2-S3).

### Optimization outcomes

All connectivity distributions (TR1, TR2, Test, Training mean, Test mean, and ΔTest) deviated from normality in both groups (Lilliefors tests, all p < 0.001), supporting the use of nonparametric inference. Mean connectivity values during TR1, TR2, and Test did not significantly differ between groups at the level of overall Test mean connectivity (Wilcoxon rank-sum p > 0.10), indicating comparable absolute connectivity strength during the final run.

In the up-regulation group, connectivity remained stable from Training to Test. Within-group comparisons revealed no significant changes from TR1 to Test (Wilcoxon signed-rank p = 0.56) or from TR2 to Test (p = 0.23). The overall Training-to-Test change score (ΔTest = Test − Training mean) was not different from zero (signed-rank p = 0.85), indicating preserved connectivity across runs. In contrast, the down-regulation group exhibited a reduction in connectivity from Training to Test. Connectivity significantly decreased from TR1 to Test (signed-rank p = 0.019), and the ΔTest change score was significantly negative (signed-rank p = 0.019), demonstrating a reliable decline in connectivity across the session.

Between-group comparison of ΔTest indicated a trend-level difference using the Wilcoxon rank-sum test (p = 0.113). However, a two-sided permutation test (20,000 label shuffles) yielded a significant group difference (p = 0.043), supporting the interpretation that connectivity was maintained in the up-regulation group but declined in the down-regulation group. Within the Test run, gain was computed as the difference between late and early connectivity (iterations 11–15 minus iterations 1–5). The between-group difference in Test gain did not reach statistical significance (rank-sum p = 0.211). Thus, group effects were more robustly observed in the Training-to-Test change metric than in within-Test connectivity dynamics.

### Working memory performance

In the Down-Regulation group, mean accuracy was 0.75 ± 0.15 at TR1, 0.77 ± 0.12 at TR2, and 0.75 ± 0.11 during the Test run, corresponding to a baseline accuracy of 0.76 ± 0.13 and a negligible change from baseline to Test (Δ = −0.004 ± 0.08). Mean reaction time (RT) was 1.08 ± 0.14 s at TR1, 0.99 ± 0.19 s at TR2, and 0.96 ± 0.20 s at Test, yielding a baseline RT of 1.04 ± 0.16 s and a modest reduction at Test (Δ = −0.07 ± 0.05 s). In the Up-Regulation group, mean accuracy increased from 0.79 ± 0.09 at TR1 and 0.82 ± 0.09 at TR2 to 0.83 ± 0.10 at Test, corresponding to a baseline accuracy of 0.81 ± 0.08 and a positive change (Δ = +0.03 ± 0.07). Mean RT values were 1.10 ± 0.28 s at TR1, 1.05 ± 0.28 s at TR2, and 0.99 ± 0.29 s at Test, with a baseline RT of 1.08 ± 0.27 s and a reduction at Test (Δ = −0.09 ± 0.10 s). Additional descriptive statistics are provided in Supplementary Table S4.

### Accuracy learning dynamics

Trial-level accuracy trajectories across TR1, TR2, and Test are illustrated in Figure 3a. During the training runs (TR1 and TR2), both groups exhibited relatively stable accuracy across binned trials, with no significant between-group differences in within-run learning slopes (all p > 0.30). In contrast, during the Test run, the Up-Regulation group demonstrated a progressive increase in accuracy across trials, whereas the Down-Regulation group showed a comparatively flatter or slightly declining trajectory. This divergence resulted in a significant between-group difference in within-run accuracy slope (p = 0.036), indicating differential accuracy learning dynamics during the Test phase.

**Figure 3.**
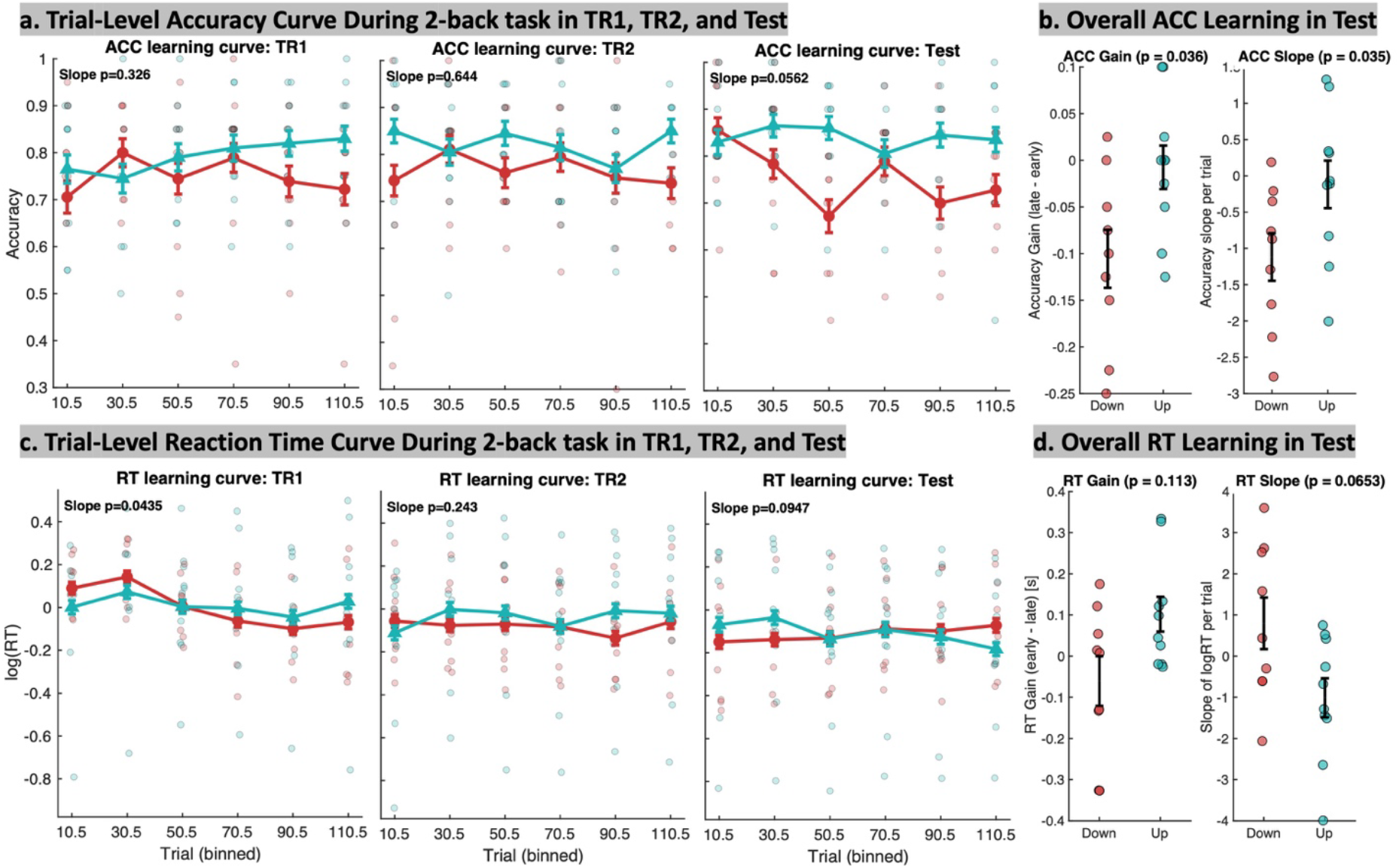
Behavioral learning dynamics during the 2-back task. (a) Trial-level accuracy learning curves across TR1, TR2, and Test. Individual participant values are shown as semi-transparent points (Down-Regulation: red; Up-Regulation: teal). Solid lines with error bars represent group mean ± standard error of the mean (SEM) across binned trials. Reported p-values indicate between-group differences in within-run accuracy learning slopes estimated from linear fits across binned trials. **(b) Test-phase accuracy learning metrics, including gain and slope**. Accuracy gain was defined as the difference between late (trials 81–120) and early (trials 1–40) accuracy within the Test run, whereas accuracy slope reflects the linear trend across individual Test trials. Points represent individual participants and error bars denote group mean ± SEM. **(c) Trial-level reaction-time (RT) learning curves (log-transformed RT) across TR1, TR2, and Test**, displayed as individual participant values with group mean ± SEM across binned trials. Reported p-values indicate between-group differences in within-run RT learning slopes. **(d) Test-phase RT learning metrics, including gain and slope**. RT gain was defined as the difference between early and late RT within the Test run (RT_early − RT_late), such that positive values reflect faster responses at the end of the run. RT slope represents the linear trend in log-transformed RT across Test trials. Points represent individual participants and error bars denote group mean ± SEM. **Abbreviations:** RT, reaction time; TR1, Training Run 1; TR2, Training Run 2; Test, Test Run; SEM, standard error of the mean.

**Figure 4.**
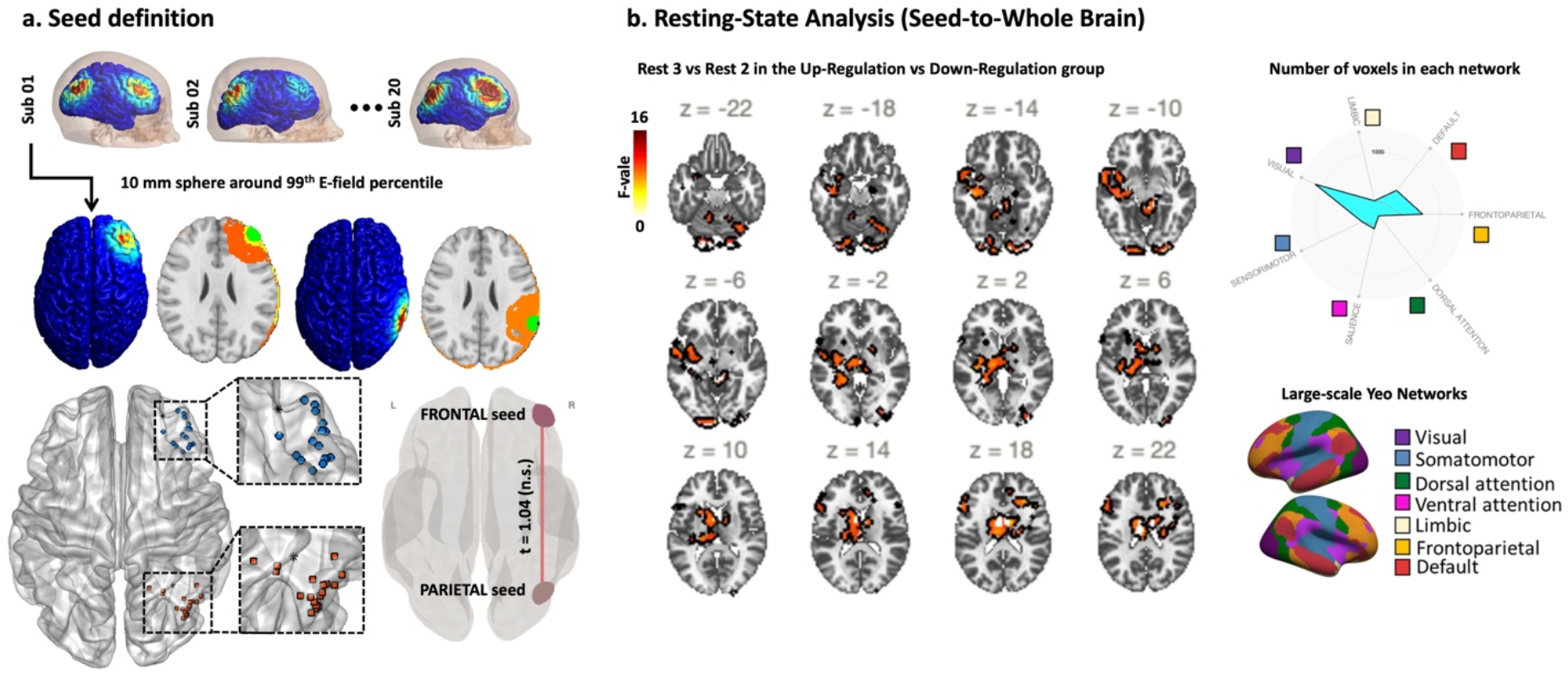
Seed definition and seed-to-whole-brain resting-state connectivity analysis. **(a)** Individualized stimulation targets were defined using participant-specific electric-field (E-field) simulations. Prefrontal and parietal seeds were derived from regions exceeding the 99th percentile of E-field magnitude and projected to MNI space to enable group-level analyses. **(b)** Seed-to-whole-brain resting-state functional connectivity results based on considering frontal and parietal seeds obtained from E-field simulations. Group-level Z-score maps are shown across representative axial slices, with warm colors indicating positive connectivity and cool colors indicating negative connectivity. Spider (radar) plot illustrating the distribution of seed-to-whole-brain connectivity across large-scale functional networks, with each axis representing a network and polygon magnitude reflecting the number of connected voxels within that network (reported in Table 2).

To further quantify within-Test performance change, accuracy gain was computed as the difference between late (trials 81–120) and early (trials 1–40) performance. The Up-Regulation group exhibited positive gain, reflecting improved accuracy across the Test run, whereas the Down-Regulation group showed minimal or negative change. Between-group comparison confirmed a significant difference in Test accuracy gain (p = 0.036; Figure 3b). Consistent with these findings, analysis of trial-wise accuracy slopes during the Test phase also revealed a significant between-group difference (p = 0.035), supporting the presence of sustained learning improvement in the Up-Regulation group. Collectively, these results indicate that group differences emerged specifically during the Test phase and were driven by distinct within-session accuracy learning dynamics rather than baseline performance differences.

### Reaction time learning dynamics

Trial-level RT trajectories (log-transformed) are shown in Figure 3c. During TR1, a significant between-group difference in within-run RT slope was observed (p = 0.043), suggesting differential early-session adaptation. However, no significant slope differences were detected during TR2 or the Test run (both p > 0.09). Across runs, both groups exhibited modest RT reductions over trials, consistent with practice-related speeding effects.

Within-Test RT gain was defined as the difference between early and late RT (RT_early − RT_late), such that positive values indicate faster responses by the end of the run. Although both groups demonstrated RT reductions across the Test phase, the between-group difference in RT gain was not statistically significant (p = 0.113; Figure 3d). Similarly, Test-phase RT slope differences did not reach significance (p = 0.065).

### Resting state connectivity

In the time-by-group interaction analysis of the seed-to-whole-brain connectivity, two significant clusters were identified when the right frontal and right parietal seeds (individualized peak electric field under the frontal/parietal electrode) were considered in the comparison between Rest 2 and Rest 3 (before and after the Test-Run as described in Figure 1) for the up-regulation versus down-regulation groups, as reported in Table 2.

## Discussion

In this study, we aimed to determine whether a real-time, closed-loop optimization framework could identify individualized stimulation parameters capable of modulating frontoparietal functional connectivity during training. We further examined whether the optimized parameters would differentially influence working memory performance (assessed using a 2-back task) and task-related connectivity during a subsequent testing phase. Finally, we investigated whether optimized stimulation produced sustained changes in intrinsic resting-state connectivity. To address these objectives, we implemented a concurrent closed-loop tES–fMRI system in which stimulation frequency and inter-site phase alignment were iteratively adjusted using an adaptive algorithm to either up-regulate or down-regulate frontoparietal synchronization during working memory engagement.

This approach yielded several key findings. First, the closed-loop optimization framework successfully identified individualized stimulation parameters aligned with real-time connectivity targets and group-specific regulation goals. Second, connectivity trajectories diverged across groups from Training to the final Test run. Whereas the Up-Regulation group maintained frontoparietal connectivity across runs, the Down-Regulation group exhibited a significant Training-to-Test decline, with the between-group difference supported by permutation testing. Third, behavioral effects were primarily observed in the accuracy domain. Despite comparable average performance levels, the Up-Regulation group demonstrated a more positive accuracy learning trajectory, greater within-Test gain, and steeper trial-wise improvement compared with the Down-Regulation group. In contrast, both groups exhibited progressive reaction-time reductions consistent with practice effects, but no robust between-group differences in Test-phase RT gain or slope were detected. Finally, resting-state analyses revealed significant time-by-group interaction effects in seed-to-whole-brain connectivity following the Test run, with connectivity changes differing across groups. Specifically, the up-regulation group exhibited increased connectivity from Rest2 to Rest3 within cerebellar and middle frontal clusters, whereas the down-regulation group showed reduced or minimal changes, indicating that optimized stimulation parameters induced directionally consistent and persistent modulation of intrinsic functional networks beyond the task context.

To close the loop between stimulation parameters and brain state, two elements are essential ^28^: (1) a brain-based marker that reflects the hypothesis and responds to stimulation, and (2) an algorithm fast enough to optimize parameters online. In our study, in line with previous frontoparietal tACS during working memory tasks ^10,35-37^, we used frontoparietal connectivity as the neural marker. We engaged this circuit with a working memory task and applied high-definition dual-site (frontal and parietal) tACS to focally target it. Our results show that the stimulation effectively engaged the frontoparietal network and that the simplex optimization method was sufficiently fast to update parameters at each iteration.

To demonstrate that the observed effects were specifically driven by the optimization algorithm, we implemented an active control condition with the opposite hypothesis about the required phase and frequency. While the up-regulation group received stimulation tailored to increase frontoparietal synchrony, the down-regulation group was stimulated to minimize it. Results showed that the up-regulation group improved working memory accuracy and slower reaction time compared to the down-regulation group, but the results did not reach a significant level.

Our results showed that in pre–post analysis, the two groups differed in terms of large-scale resting-state network connectivity. The main network that demonstrated significantly higher connectivity to the stimulation site was the frontoparietal network. tES-induced resting-state functional connectivity changes between stimulation site and large-scale brain networks, including FPN and default mode networks, after tACS and working memory have been reported in previous studies^38-40^.

However, consistent with prior pre–post tACS resting-state fMRI studies^38^, changes in offline resting-state connectivity were not directly associated with online behavioral improvements. This dissociation likely reflects the state-dependent nature of the stimulation protocol. In the present study, stimulation parameters were optimized in real time during active working memory engagement to modulate task-evoked frontoparietal synchronization. As such, the identified frequency and phase parameters were tuned to influence network dynamics under cognitive load, rather than intrinsic connectivity in the absence of task demands. Online behavioral effects are therefore likely driven by transient, phase-specific modulation of interregional coupling during task performance. In contrast, resting-state fMRI captures spontaneous low-frequency BOLD fluctuations and reflects a more global and temporally averaged estimate of intrinsic network organization. The mechanisms underlying acute oscillatory entrainment during task performance may not necessarily translate into durable plastic changes detectable during subsequent rest.

Although we considered a large search space, the final phase and frequency values for each group did not fall into clearly distinct ranges (e.g., in-phase vs. anti-phase, or low theta vs. high theta). This finding highlights two important points. First, due to safety considerations, we limited stimulation in intensity and duration, which restricted the strength and number of optimization iterations; more iterations may be necessary to identify the ultimate optimal parameters. Second, the relatively narrow variability in frequency ranges across groups raises questions about the necessity of strict frequency/phase individualization, at least in healthy adults (as suggested before in comparison between closed vs open loop tACS ^26^ or a low range of standard deviation in the individualized frequency reported before^38^). A standardized frequency might be sufficient, potentially simplifying tACS protocols without reducing efficacy. To address these points, future studies should incorporate longer stimulation durations, a larger number of optimization iterations, and sham-controlled comparisons of peak, trough, and random phase/frequency conditions, or employ predefined search spaces for each condition.

Additionally, electric field modeling offers a complementary approach for protocol refinement. Specifically, greater overlap between the tES-induced electric field and the fMRI-derived network-level pattern was significantly associated with the outcome ^41^. Furthermore, scalp-to-cortex distance has been shown to create significant differences in HD-tACS–induced electric field strength between frontal and parietal sites, and the symmetry of the induced electrical field magnitudes influenced the task performance during various stimulation conditions^17,18,42^. Achieving balanced fields, therefore, requires lower stimulation intensity over parietal regions. Second, a recent open-label clinical trial in 10 participants with major depressive disorder implemented a bifrontal, individualized alpha frequency–based closed-loop tACS-EEG protocol with duration tailored to each patient’s ongoing brain activity^43^. Specifically, stimulation was triggered whenever power in the individual alpha frequency (IAF) band—estimated from 10-s EEG segments—exceeded a predefined threshold, ensuring alignment with peaks of alpha activity rather than periods of low power^43^. This approach produced significant improvements in mood outcomes (p < 0.001; 80% response and remission rate), underscoring the value of individualized timing in optimizing stimulation dose and efficacy^43^. Taken together, these findings suggest that future protocols may benefit from balancing both stimulation intensity (to account for individual anatomy) and stimulation duration (to account for ongoing brain state) ^44^.

Although our study provides promising preliminary evidence from the first closed-loop tACS-fMRI investigation in a group of participants, several limitations should be acknowledged. First, the relatively small sample size limits the generalizability of our findings and calls for replication in larger cohorts. Second, we examined only a single session of low-intensity stimulation with a restricted duration, which also constrained the number of optimization iterations. Future research should explore the effects of repeated tACS sessions, evaluate the test–retest reliability of individual responsiveness, and consider systematic comparisons between peak, trough, and random phase/frequency conditions within predefined search spaces.

Together, these results support the feasibility of closed-loop, individualized tACS-fMRI for enhancing cognitive control networks and highlight the potential importance of tailoring both stimulation parameters and timing to individual brain dynamics.

## Data and Code Availability

The data and code that support the findings of this study will be available from the corresponding author upon request after publication.

