## Supplementary Materials for "Closing the loop between brain and electrical stimulation: A proof-of-concept randomized trial of real-time fMRI-guided tACS optimization"

### S1. CONSORT Diagram

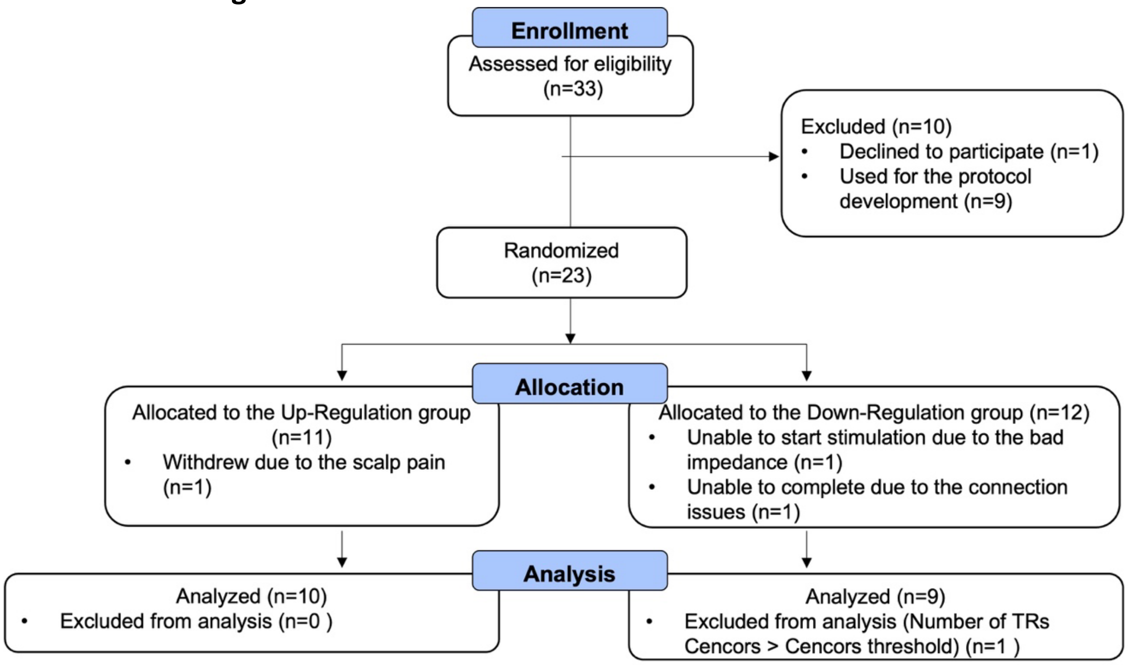


**Figure S1. CONSORT flow diagram for the study.** n: number of participants. Experimental group: a group of participants that received optimized tACS parameters to induce the highest frontoparietal connectivity. Control group: a group of participants that received controlled tACS parameters to induce the lowest frontoparietal connectivity.

### S2. Profile Of Mood State (POMS) questionnaire

The Profile of Mood States (POMS) is a validated self-report questionnaire used to assess transient mood states over a specified time frame. It consists of 65 mood-related adjectives rated on a 5-point Likert scale ranging from 0 (*not at all*) to 4 (*extremely*). POMS yields six subscale scores—Tension–Anxiety, Depression–Dejection, Anger–Hostility, Vigor–Activity, Fatigue–Inertia, and Confusion–Bewilderment—and a global index of mood, the Total Mood Disturbance (TMD), calculated by summing all negative mood subscales and subtracting Vigor. The instrument is widely used in clinical and experimental research and is particularly sensitive to short-term mood changes, making it well-suited for repeated-measures designs and intervention studies.

**Table S1. POMS.** Between-group comparison

| | Measure | N_down | Mean_down | SD_down | N_up | Mean_up | SD_up | t | df | p_value |
|---|---|---|---|---|---|---|---|---|---|---|
| pre | poms_tension_score | 9 | 2.667 | 2.062 | 10 | 3.300 | 2.946 | -0.547 | 16.098 | 0.592 |
| pre | poms_depression_score | 9 | 0.222 | 0.441 | 10 | 1.900 | 4.977 | -1.061 | 9.157 | 0.316 |
| pre | poms_anger_score | 9 | 1.000 | 1.658 | 10 | 1.200 | 2.821 | -0.191 | 14.787 | 0.851 |
| pre | poms_fatigue_score | 9 | 1.556 | 2.007 | 10 | 2.000 | 2.160 | -0.465 | 16.976 | 0.648 |
| pre | poms_confusion_score | 9 | 3.111 | 0.782 | 10 | 2.800 | 2.530 | 0.370 | 10.873 | 0.719 |
| pre | poms_vigour_score | 9 | 13.667 | 7.246 | 10 | 18.100 | 6.773 | -1.373 | 16.474 | 0.188 |
| pre | poms_tmd_score | 9 | -5.111 | 9.075 | 10 | -6.900 | 17.496 | 0.284 | 13.798 | 0.781 |
| post | poms_tension_score | 8 | 4.000 | 3.117 | 8 | 3.250 | 2.765 | 0.509 | 13.803 | 0.619 |
| post | poms_depression_score | 8 | 0.250 | 0.463 | 8 | 1.250 | 3.536 | -0.793 | 7.240 | 0.453 |
| post | poms_anger_score | 8 | 1.000 | 1.512 | 8 | 1.250 | 2.816 | -0.221 | 10.726 | 0.829 |
| post | poms_fatigue_score | 8 | 2.375 | 1.685 | 8 | 4.250 | 3.694 | -1.306 | 9.793 | 0.221 |
| post | poms_confusion_score | 8 | 2.875 | 0.835 | 8 | 3.375 | 2.326 | -0.572 | 8.773 | 0.582 |
| post | poms_vigour_score | 8 | 11.500 | 6.141 | 8 | 17.125 | 7.140 | -1.689 | 13.694 | 0.114 |
| post | poms_tmd_score | 8 | -1.000 | 11.464 | 8 | -3.750 | 16.977 | 0.380 | 12.285 | 0.711 |

**Table S2. POMS.** Pre-Post comparison

| Group | Measure | N_pairs | Mean_pre | Mean_post | t | df | p_value |
|---|---|---|---|---|---|---|---|
| down-regulation | poms_tension_score | 8 | 2.5 | 4 | -1.080 | 7 | 0.316 |
| down-regulation | poms_depression_score | 8 | 0.25 | 0.25 | 0.000 | 7 | 1.000 |
| down-regulation | poms_anger_score | 8 | 0.875 | 1 | -0.196 | 7 | 0.850 |
| down-regulation | poms_fatigue_score | 8 | 1.75 | 2.375 | -0.691 | 7 | 0.512 |
| down-regulation | poms_confusion_score | 8 | 3 | 2.875 | 0.552 | 7 | 0.598 |
| down-regulation | poms_vigour_score | 8 | 13 | 11.5 | 1.426 | 7 | 0.197 |
| down-regulation | poms_tmd_score | 8 | -4.625 | -1 | -1.426 | 7 | 0.197 |
| up-regulation | poms_tension_score | 8 | 3.25 | 3.25 | 0.000 | 7 | 1.000 |
| up-regulation | poms_depression_score | 8 | 2.25 | 1.25 | 1.366 | 7 | 0.214 |
| up-regulation | poms_anger_score | 8 | 1.375 | 1.25 | 0.314 | 7 | 0.763 |
| up-regulation | poms_fatigue_score | 8 | 2.5 | 4.25 | -1.941 | 7 | 0.093 |
| up-regulation | poms_confusion_score | 8 | 2.75 | 3.375 | -1.667 | 7 | 0.140 |
| up-regulation | poms_vigour_score | 8 | 18.375 | 17.125 | 0.798 | 7 | 0.451 |
| up-regulation | poms_tmd_score | 8 | -6.25 | -3.75 | -0.798 | 7 | 0.451 |

### S3. State–Trait Anxiety Inventory—State form (STAI-S)

The State–Trait Anxiety Inventory—State form (STAI-S) is a widely used, validated self-report measure designed to assess current (state) anxiety, reflecting how individuals feel "right now". The STAI-S consists of 20 items, each rated on a 4-point Likert scale ranging from 1 (*Not at all*) to 4 (*Very much so*). Items capture both anxiety-present (e.g., feeling tense, nervous, worried) and anxiety-absent states (e.g., feeling calm, relaxed, secure). To compute the State Anxiety score, anxiety-absent items are reverse-scored and summed with anxiety-present items, yielding a total score in which higher values indicate greater state anxiety.

**Table S3. STAI-S.** Between-group comparison

| Group | Question | N_pairs | Mean_pre | Mean_post | t | df | p_value |
|---|---|---|---|---|---|---|---|
| down-regulation | STAI-State Total Score | 8 | 30.750 | 33.250 | -0.960 | 7 | 0.369 |
| up-regulation | STAI-State Total Score | 9 | 28.333 | 26.111 | 0.845 | 8 | 0.422 |

**Table S4. STAI-S.** Pre-Post comparison

| Session | Question | N_down | Mean_down | SD_down | N_up | Mean_up | SD_up | t | df | p_value |
|---|---|---|---|---|---|---|---|---|---|---|
| pre | STAI-State Total Score | 9 | 30.667 | 7.159 | 10 | 28.600 | 6.450 | 0.658 | 16.250 | 0.520 |
| post | STAI-State Total Score | 8 | 33.250 | 8.828 | 9 | 26.111 | 11.494 | 1.445 | 14.728 | 0.169 |

### S4. More results on behavioral performance

Linear mixed-effects (LME) models with Run and Group as fixed effects (subject-level random effects; residual degrees of freedom) showed no significant main effects or interactions for accuracy, including no effect of Group ($F(1,51) = 0.84$, $p = 0.36$), Run ($F(2,51) = 0.25$, $p = 0.78$), or Group × Run interaction ($F(2,51) = 0.58$, $p = 0.57$). In contrast, reaction time showed a significant main effect of Run ($F(2,51) = 6.88$, $p = 0.002$), indicating changes in RT across phases, while neither the main effect of Group ($F(1,51) = 0.05$, $p = 0.83$) nor the Group × Run interaction ($F(2,51) = 0.45$, $p = 0.64$) reached significance.

Planned contrasts examining within-group changes from Baseline to Test showed no significant change in accuracy in either group (Down-Regulation: $t(8) = -0.15$, $p = 0.886$; Up-Regulation: $t(9) = 1.20$, $p = 0.259$). In contrast, reaction time decreased significantly from Baseline to Test in both groups (Down-Regulation: $t(8) = -4.11$, $p = 0.003$; Up-Regulation: $t(9) = -2.94$, $p = 0.017$). Planned between-group contrasts on change scores (Δ Test–Baseline) revealed no significant group differences for either accuracy ($t(16.2) = -0.89$, $p = 0.387$) or reaction time ($t(14.3) = 0.47$, $p = 0.646$).

Across models, beliefs about receiving real stimulation and confidence were not associated with accuracy change (all $p > 0.17$). Side-effect burden showed a weak trend toward a positive association with accuracy change ($\beta \approx 0.02$), but this effect did not reach statistical significance and was not robust across models. No significant Group × Moderator interactions were observed for age, sex, baseline working memory performance, impedance, E-field strength, or head motion (all $p > 0.10$).
